# Yet another mitochondrial genome of the Pacific cupped oyster: the published mitogenome of *Alectryonella plicatula* (Ostreinae) is based on a misidentified *Magallana gigas* (Crassostreinae)

**DOI:** 10.1101/2021.06.19.449104

**Authors:** Daniele Salvi, Emanuele Berrilli, Matteo Garzia, Paolo Mariottini

**Affiliations:** Department of Health, Life and Environmental Sciences, University of L’Aquila, Via Vetoio, 67100 Coppito, L’Aquila, Italy; Dipartimento di Scienze, Università Roma Tre, Viale G. Marconi 446, 00146 Rome, Italy

**Author notes:** Corresponding author: Daniele Salvi.

**Keywords:** DNA barcoding, *Magallana*, misidentification, Ostreidae, oyster, phylogeny

## Abstract

The recently published mitochondrial genome of the fingerprint oyster *Alectryonella plicatula* (Gmelin, 1791) with GenBank accession number MW143047 was resolved in an unexpected phylogenetic position, as sister to the Pacific cupped oyster *Magallana gigas* (Thunberg, 1793) and share with this species three typical gene duplications that represent robust synapomorphies of the *Magallana* clade. In this study, we verified the identity of MW143047 using direct comparisons of single gene sequences, DNA barcoding and phylogenetic analyses. BLAST searches using each of the 12 protein coding genes and rRNA genes extracted from MW143047 as query retrieved *M. gigas* as best hit with 100% sequence identity. MW143047 is nested within the clade formed by *M. gigas* sequences, with virtually no difference between their terminal branch lengths, both in the *cox1* gene tree (based on 3639 sequences) and in the 16S gene tree (based on 1839 sequences), as well as in the Maximum Likelihood mitogenomic tree based on concatenated sequence of 12 PCGs. Our findings suggest that the original specimen used for mitogenome sequencing was misidentified and represents an individual of *M. gigas*. This study reinforces the notion that morphological shell analysis alone is not sufficient for oyster identification, not even at high taxonomic ranks such as subfamilies. While it is well established that morphological identification of oysters should be validated by molecular data, this study emphasizes that also molecular data should be taxonomically validated by means of DNA barcoding and phylogenetic analyses. The implications of the publication of taxonomically misidentified sequences and mitogenomes are discussed.

## Introduction

Oysters are distributed worldwide in temperate and tropical waters and several of them have a great economic importance. However, taxonomic identification of oysters based on morphological characters is challenging, even for species locally cultivated since centuries (e.g. Wang et al., 2004; Hsiao et al., 2016). Indeed, oysters’ shells show a high degree of phenotypic plasticity driven by environmental factors, therefore, shell morphology is often uninformative or misleading for taxonomic identification and classification. The use of molecular data has been fruitful for species identification and has resulted in a well-established phylogeny and systematics of oysters (Salvi et al., 2014; Salvi & Mariottini, 2017). The mitochondrial genome has been the most valuable source of molecular data for oyster species identification (DNA barcoding), phylogenetic reconstruction and classification (e.g. Wang et al., 2004; Liu et al., 2011; Salvi et al., 2014; Raith et al., 2016). Moreover, mitochondrial gene rearrangements, such as transpositions and duplications, has provided additional characters for phylogenetic inference, classification and diagnosis of oysters’ genera and subfamilies (Salvi & Mariottini, 2021). Molecular resources of oyster are continuously growing, and most studies currently implement these data for taxonomic identification. For this purpose, a reliable reference of taxonomically identified sequences and mitogenomes is necessary (Bortolus, 2008; Jin et al., 2020; Salvi et al., 2020).

Recently, the complete mitochondrial genome of the fingerprint oyster *Alectryonella plicatula* (Gmelin, 1791), with GenBank accession number MW143047, has been characterised (Wang et al., 2021) and resolved in an unexpected phylogenetic position, as sister to the Pacific cupped oyster *Magallana gigas* (Thunberg, 1793). Unfortunately, in this mitogenome announcement the phylogenetic position of MW143047 is described in a cladogram with arbitrary branch lengths (Wang et al., 2021), therefore masking the true evolutionary divergence between MW143047 and the mitogenome of *M. gigas* (see Botero-Castro et al., 2016). However, their sister relationship is surprising and in sharp contrast with all previous phylogenetic studies that have consistently established the placement of *A. plicatula* within the lophinae lineage, that is nested within the subfamily Ostreinae Rafinesque, 1815, whereas *M. gigas* belong to the well-defined clade of Indo-Pacific Crassostreinae Scarlato & Starobogatov, 1979 (O’Foighil & Taylor, 2000; Salvi et al., 2014; Crocetta et al., 2015; Salvi & Mariottini, 2017; Al-Kandari et al., 2021). Moreover, the newly published mitogenome MW143047 conforms to the mitochondrial gene arrangement of *M. gigas*, that is characterised by the duplication of *trnK, trnQ* and *rrnS* genes that are exclusive of the *Magallana* clade (Ren et al., 2010) and represent robust synapomorphies of this clade (Salvi et al., 2014; Salvi & Mariottini, 2017, 2021). These intriguing points are urgent to clarify as MW143047 might become the mitogenomic reference of *A. plicatula*. In this study, we verified the taxonomic identification of Wang et al (2021) using available quality control guidelines for taxonomic validation of new mitogenomes (Botero-Castro et al., 2016).

## Materials and Methods

We verified the identity of MW143047 using DNA barcoding and phylogenetic analyses.

We extracted from the mitogenome MW143047 the two barcoding fragments commonly used for oysters, the *cox1* and the 3’ half portion of the 16S rRNA (Liu et al 2011; Crocetta et al., 2015), as well the remaining protein coding genes and rRNAs (12S and the 5’ half portion of the 16S) using Geneious Prime 2021 (Biomatters Ltd., Auckland, New Zealand). Sequence of each gene were used as query in BLAST searches using default settings. Sequences of the barcoding markers *cox1* and the 16S were aligned with oysters’ sequences available from public database (BOLD and NCBI) assembled, dereplicated, and aligned following the procedure by Salvi et al. (2020). A Neighbor-Joining (NJ) tree was constructed based on uncorrected *p*-distance values in MEGA v. 7 (Kumar et al., 2016) with pairwise deletion and 100 replicates of bootstrap (BS).

We inferred a Maximum Likelihood (ML) tree based on the concatenated sequences of 12 protein-coding genes (PCGs) of the same oyster taxa analysed by Wang et al. (2021) plus six additional mitogenome sequences of *M. gigas*, to further assess phylogenetic relationships and divergence between the latter and the mitogenome MW143047. ML analyses were performed in IQTREE v 1.6.12 (Nguyen et al., 2015) using for each gene partition the best substitution model determined by the ModelFinder module (Kalyaanamoorthy et al., 2017) and 1000 replicates of ultrafast bootstrapping.

## Results

Results of BLAST searches using as query the *cox1* and the 16S sequences extracted from MW143047 retrieved as best hits sequences assigned to *M. gigas* with a sequence identity of 100% (sequence identity ranging from 99.85% to 100% among the best 10 hits for *cox1* and of 100% for 16S; Table 1). The same result was obtained in BLAST searches using as query the other 11 protein coding genes and rRNAs extracted from MW143047, with 100% of nucleotides identical to multiple sequences of *M. gigas*.

**Table 1.**
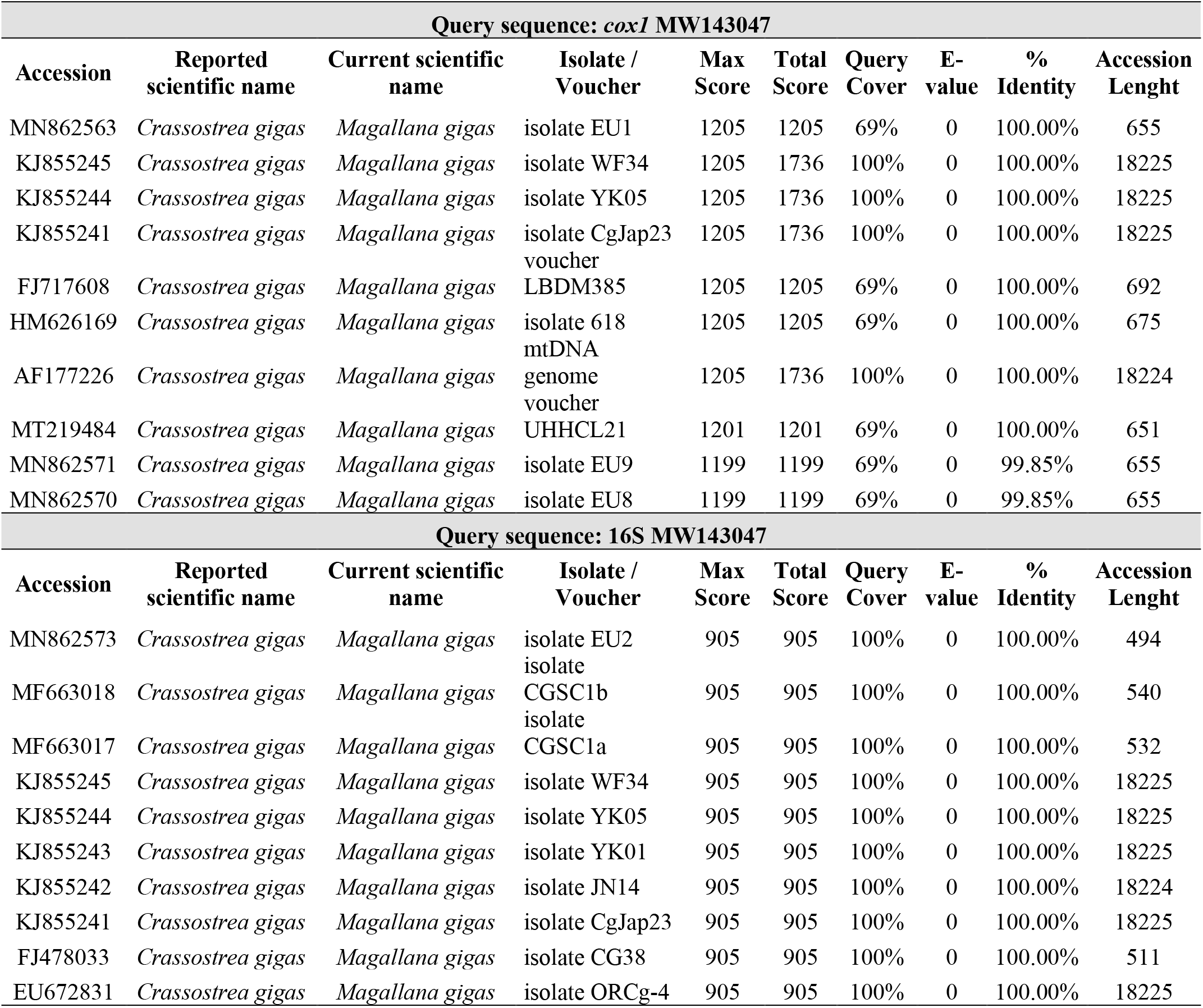
Top ten best hits of BLAST results using as query the sequences of the barcoding fragments *cox1* (above) and 16S rRNA (below) extracted from the complete mitochondrial genome MW143047.

In the gene tree based on 3639 *cox1* sequences (Fig 1a) and in the gene tree based on 1839 16S sequences (Fig 1b) MW143047 clustered with *M. gigas* with maximum bootstrap support (BS=100%).

**Figure 1.**
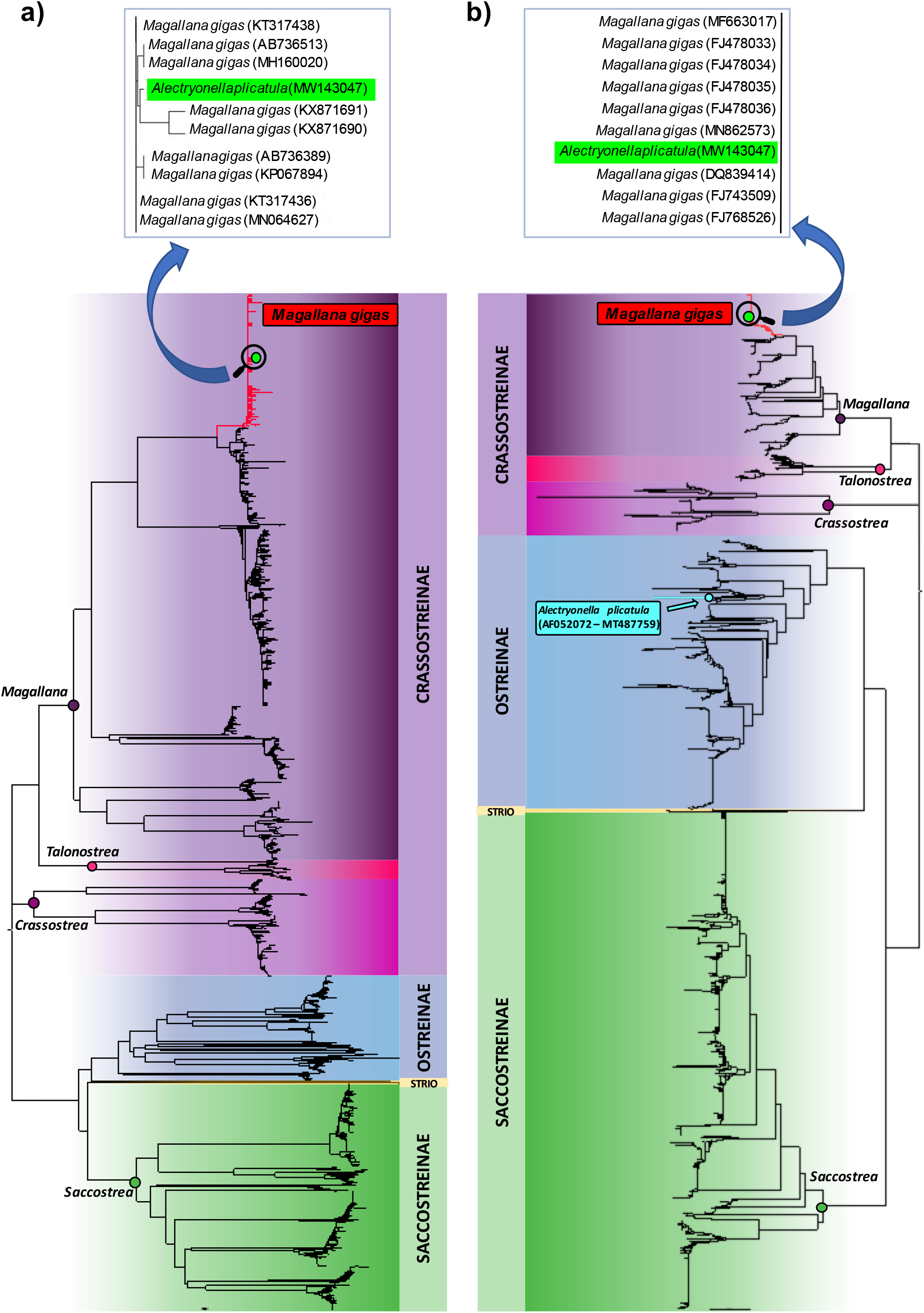
Neighbor-Joining trees based on 3639 *cox1* sequences (a) and 1839 16S sequences (b) available from public databases. In both trees MW143047 is nested within the clade formed by sequences of *Magallana gigas* within the Crassostreinae lineage. Instead, available 16S rRNA sequences of *Alectryonella plicatula* generated in previous studies cluster within the Ostreinae lineage. (STRIO:Striostreinae).

In the ML mitogenomic tree (Fig 2) MW143047 is nested within the clade formed by *M. gigas* sequences, with virtually no difference between their terminal branch lengths. This clade was sister to the mitogenome sequence of *M. angulata* (BS=100%) within the well supported clade formed by *Magallana* species (BS=100%).

**Figure 2.**
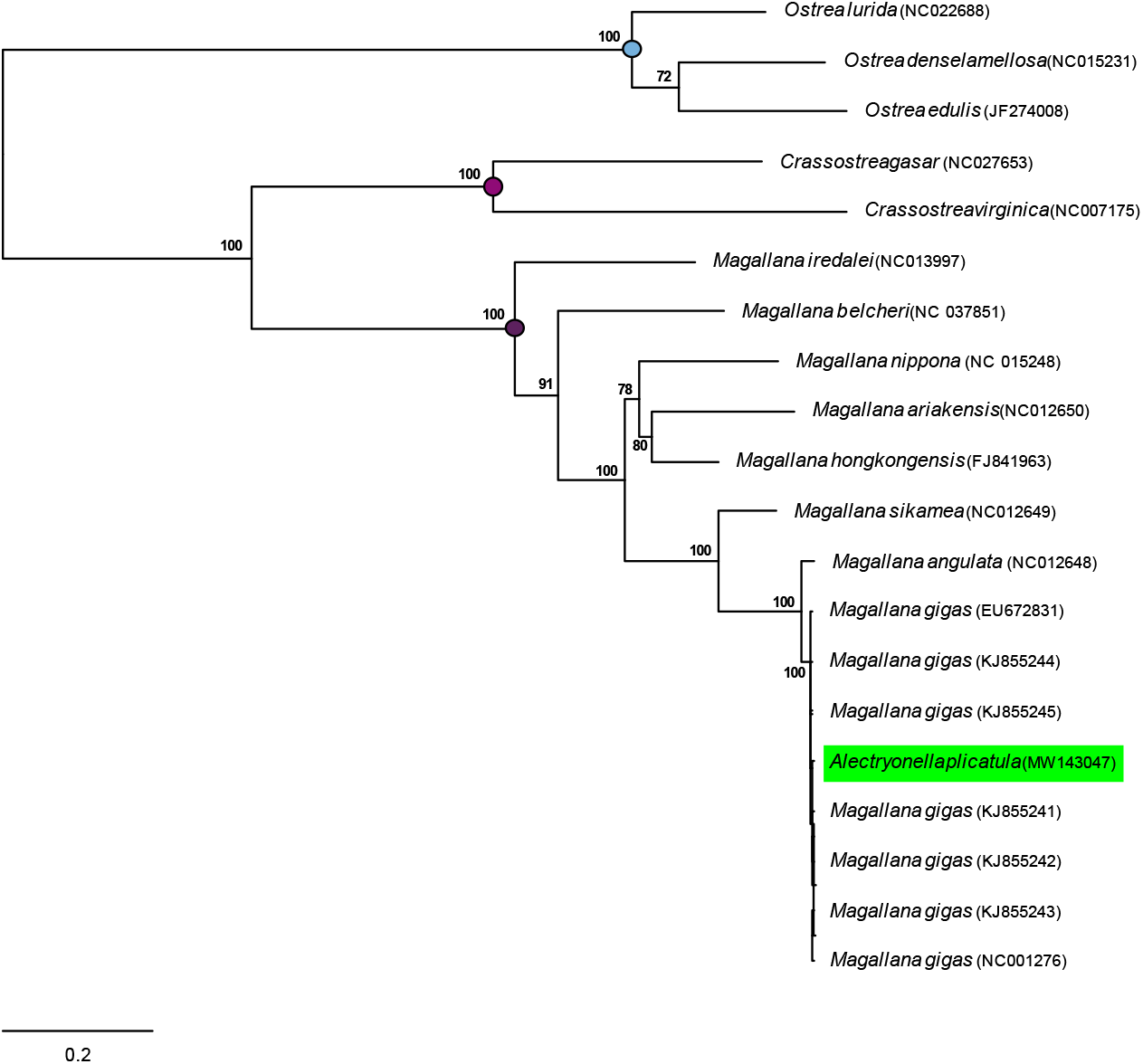
Maximum likelihood tree based on the concatenated sequences of 12 protein-coding from complete mitochondrial genomes of the same oyster taxa analyse d by Wang et al. (2021) plus six additional mitogenome sequences of *Magallana gigas*. The mitogenome MW143047 is nested with the clade formed by mitogenomes of *M. gigas*.

## Discussion

Results of DNA barcoding, BLAST and phylogenetic analyses show that MW143047, attributed by Wang et (2021) to the fingerprint oyster *Alectryonella plicatula*, is identical to mitochondrial DNA sequences of the Pacific cupped oyster *M. gigas* (Table 1). The MW143047 sequences cluster within the clade of *M. gigas* both in the gene trees based on the barcoding markers *cox1* and 16S and in the ML mitogenome tree based on concatenated sequence of 12 PCGs (Fig 1 and 2). On the other hand, two mitochondrial 16S rRNA sequences of *A. plicatula* generated in previous studies (Jozefowicz & O’Foighil, 1998; Ardura et al., 2021), and available in Genbank under the accession numbers (AF052072 and MT487759), show a high genetic divergence (*p*-distance: 19 and 18% respectively) with MW143047. The most likely explanation for these results is that the original specimen used for mitogenome sequencing was misidentified and represents an individual of *M. gigas*.

The hypothesis of contamination by DNA of *M. gigas*, either prior to PCR amplification or as PCR product prior to sequencing, is unlikely. In these cases, often chimera sequence artefacts are observed (e.g., Sangster & Luksenburg, 2020), whereas all PCGs and rRNA genes of MW143047 are identical to sequences of *M. gigas* thus indicating that MW143047 is a *bona fide* mitogenome of *M. gigas*. Even less likely is the hypothesis of mitochondrial introgression of *M. gigas* in *A. plicatula* following hybridization. Indeed, while these two species might co-occur in the collection site of the original specimen used for sequencing (Shicheng Island, Dalian; China), their genetic divergence is very large (∼19% at the 16S rRNA) as they belong to distinct evolutionary lineages within Ostreidae Rafinesque, 1815 (*A. plicatula* belongs to the Ostreinae lineage whereas *M. gigas* to the Crassostreinae lineage; e.g. O’Foighil & Taylor, 2000; Salvi et al., 2014; Crocetta et al., 2015; Salvi & Mariottini, 2017; Al-Kandari et al., 2021).

While *Magallana gigas* in *Alectryonella plicatula* are readily distinguishable using mitochondrial (Liu et al., 2011; Crocetta et al., 2015) or nuclear markers (O’Foighil & Taylor, 2000; Salvi et al., 2014; Mazón-Suástegui et al., 2016), morphological misidentification between the two might be easy as reported by Bishop et al (2017) due the extensive degree of phenotypic plasticity of oysters. This example highlights the common difficulties encountered for identifying oysters based on shell morphology alone, and provides one more demonstration that misidentification regards not only closely related species but also taxonomic ranks as high as subfamilies (discussed in Salvi & Mariottini, 2021; see Salvi et al. 2014 and Raith et al. 2016 for examples regarding the subfamilies Striostreinae Harry, 1985, Ostreinae Rafinesque, 1815 and Saccostreinae Salvi & Mariottini, 2016).

Previous studies on oyster systematics strongly advice that morphological identification of oysters should be validated by molecular data (e.g. Wang et al., 2004; Lam & Morton, 2006; Hamaguchi et al., 2107). This study also emphasizes that molecular data should be taxonomically validated by means of DNA barcoding and phylogenetic analyses. Taxonomic validation of mitogenomes is straightforward following the quality control guidelines of Botero-Castro et al. (2016) (see also Sangster & Luksenburg, 2020) and most of these recommendations can be applied also for an accurate identification of the sequences of single gene fragments. The publication of taxonomically misidentified sequences and mitogenomes can have profound implications if few sequences are available for the species so that misidentified sequence ends up as the reference for the species in public databases. In such cases misidentification errors can propagate in future studies that use the wrong reference-sequences in taxonomic and phylogenetic comparisons.

## Data availability statement

The data that support the findings of this study are openly available on GenBank at https://www.ncbi.nlm.nih.gov/nucleotide. Accession numbers of mitogenome sequences analysed are listed in Figure 2. Results of Blast and DNA-barcoding analyses are available from the authors upon request.

